# Lungs contribute to solving the frog’s cocktail party problem by enhancing the spectral contrast of conspecific vocal signals

**DOI:** 10.1101/2020.06.30.171991

**Authors:** N. Lee, J. Christensen-Dalsgaard, L. A. White, K. M. Schrode, M. A. Bee

## Abstract

Noise impairs signal perception and is a major source of selection on animal communication. Identifying adaptations that enable receivers to cope with noise is critical to discovering how animal sensory and communication systems evolve. We integrated biophysical and bioacoustic measurements with physiological modeling to demonstrate that the lungs of frogs serve a heretofore unknown noise-control function in vocal communication. Lung resonance enhances the signal-to-noise ratio for communication by selectively reducing the tympanum’s sensitivity at critical frequencies where the tuning of two inner ear organs overlaps. Social network analysis of citizen-science data on frog calling behavior indicates the calls of other frog species in multi-species choruses are a prominent source of environmental noise attenuated by the lungs. These data reveal that an ancient adaptation for detecting sound via the lungs has been evolutionarily co-opted to create spectral contrast enhancement that contributes to solving a multi-species cocktail party problem.

## INTRODUCTION

Environmental noise is a major source of selection that shapes the evolution of animal sensory and communication systems (*1, 2*). Noise creates problems for effective acoustic communication by increasing signal detection thresholds, disrupting source localization and sound pattern recognition, and impairing auditory discrimination. There is widespread and increasing interest in understanding how animals are adapted to cope with noise problems (*1, 2*), particularly in light of the recent global rise in anthropogenic noise pollution (*3*). For many insects (*4*), frogs (*5*), and birds (*6*) that signal acoustically in large and often multi-species aggregations, the signals of other individuals represent particularly potent sources of environmental noise that reduce the signal-to-noise ratio for communication. In essence, such species must solve multi-species analogs of the human “cocktail party problem,” which refers to our difficulty communicating with speech in noisy crowds (*7, 8*). At present, knowledge of the potential diversity of adaptations for solving noise problems among nonhuman animals remains limited.

Frogs represent key taxa for discovering evolved solutions to noise problems (*5, 9, 10*). Among terrestrial vertebrates, acoustic communication evolved independently in frogs some 200 million years ago (*11*), probably soon after the independent evolution during the Triassic of a tympanic middle ear in the lineage leading to modern frogs (*12*). Vocal communication is fundamental to reproduction in most extant frogs. Male frogs aggregate in dense breeding “choruses,” often comprising multiple species, where they produce sexual signals termed “advertisement calls” to attract mates and repel rivals (*5, 9, 10*). Advertisement calls are produced at notoriously high sound amplitudes, with peak sound pressure levels (SPL) exceeding 100 dB (at 1 m) in many species (*9*). Consequently, frog choruses are often characterized by high and sustained levels of background noise that may be audible to humans from distances of up to 2 km (*5*). Within the cacophony of a chorus, female frogs must select and locate a mate based on evaluating the advertisement calls of different males (*9*). Chorus noise and the overlapping calls of conspecific and heterospecific males can negatively impact auditory perception and degrade the acoustically-guided mating decisions of females (*5*).

Amphibians are unique among vertebrates in having inner ears with two physically distinct sensory papillae that transduce different frequency ranges of airborne sounds (*9*). In frogs, the two organs are considered “matched filters” in the spectral domain because they are most sensitive to one or more spectral components in conspecific advertisement calls (*9*). The tympanic middle ears of frogs are internally coupled through the mouth cavity via wide and open Eustachian tubes (Fig. 1A) (*13, 14*). Thus, each tympanum’s response reflects the summation of sound impinging directly on its external surface and sound reaching its internal surface via input through the opposite, internally coupled tympanum (*13*). In frogs, sound can also reach the internal surface of each tympanum through the body wall and air-filled lungs via the glottis, mouth, and Eustachian tubes (Fig. 1B) (*15-19*). Prior to the evolutionary origins of tympanic middle ears, sound induced vibration of air-filled lungs was a likely mechanism of sound reception in the earliest amphibian ancestors of all tetrapods (*20*). Among extant terrestrial vertebrates, however, the lung-to-ear sound transmission pathway is unique to amphibians. During the respiratory cycle, frog lungs remain continuously pressurized above atmospheric pressure, and they remain inflated for relatively long periods punctuated by brief episodes of ventilation when pulmonary air is expelled through the glottis and then refilled using an active pump mechanism driven by buccal musculature (*18, 21*). Thus, during normal respiration, there is a strong coupling between the lungs and the air-filled tympanic middle ears of frogs. Existing hypotheses for the function of the lung-to-ear pathway include protecting a male frog’s hearing during vocalization (*14*) and sharpening the inherent directionality of their internally coupled ears (*15-19*); however, these functions are not yet well understood, and the hypothesis that the lungs improve localization of conspecific signals remains controversial (*13, 22*).

**Fig. 1.**
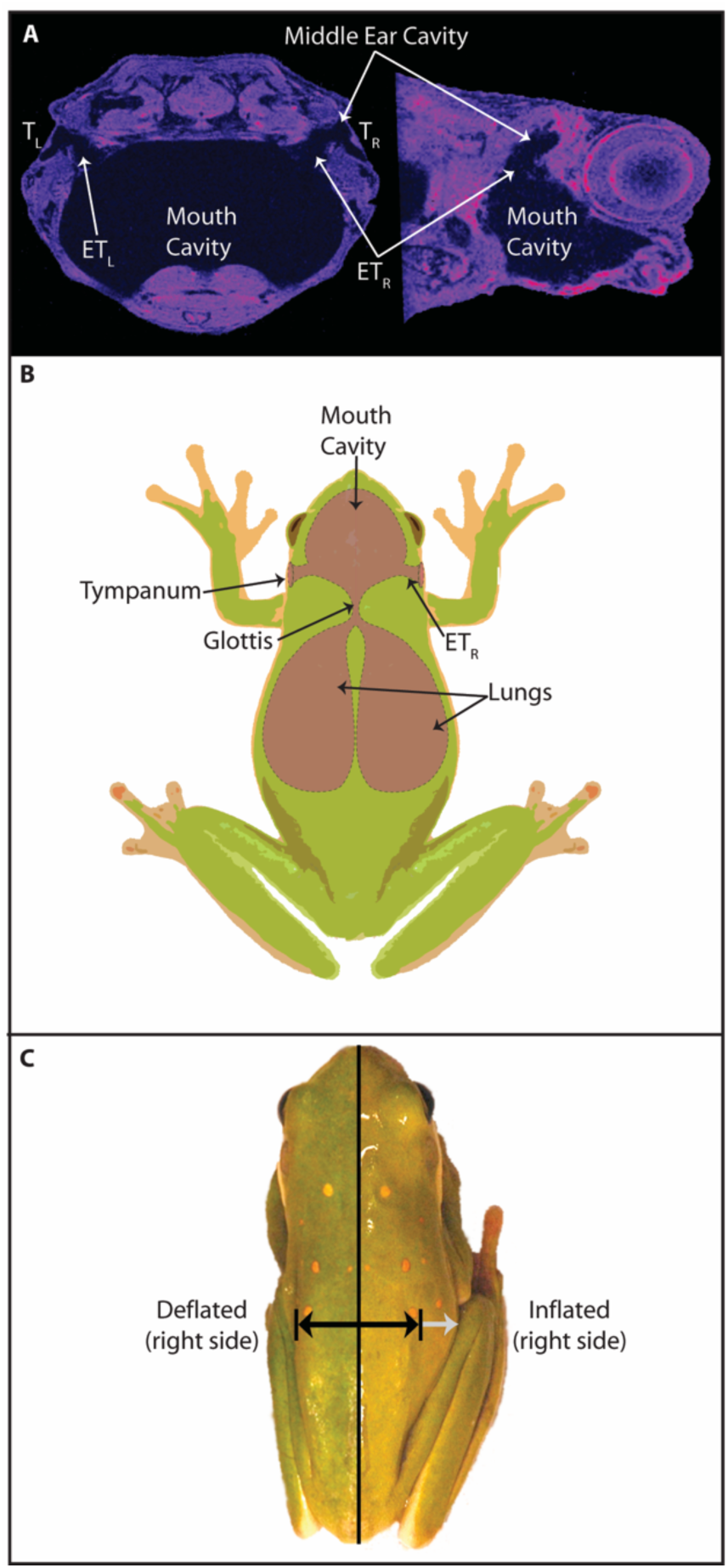
Sound transmitted via the lungs reaches the internal surfaces of the frog’s internally coupled tympana. **(A)** Magnetic resonance images (coronal, left; sagittal, right) of a green treefrog head showing the internal coupling of the left (L) and right (R) tympana (T) and air-filled middle ear cavities through the Eustachian tubes (ET) and mouth cavity. **(B)** Schematic illustration showing the coupling of the lungs through the glottis to the internally coupled tympana. **(C)** A split image of a female green treefrog showing the lateral extension of her right body wall in the inflated (non-reflected right half of image) and deflated (reflected left half of image) states of lung inflation. Black arrows depict the lateral extension of the female’s right body wall in the deflated state, and the light gray arrow depicts the additional lateral extension of the right body wall in the inflated condition.

Here, we tested a novel hypothesis for the function of the frog’s lung-to-ear sound transmission pathway: frog’s lungs improve the signal-to-noise ratio for vocal communication by creating *spectral contrast enhancement* (SCE). Spectral contrast refers to the difference in amplitude (in dB) between the “peaks” and “valleys” in a sound spectrum. In the context of human hearing and speech communication, signal processing algorithms for SCE can be used to amplify the formant frequencies (peaks) of voiced speech sounds or attenuate frequencies between adjacent formants (valleys) (*23-25*). SCE can improve speech perception in noise for hearing impaired listeners when implemented as part of the signal processing strategies of hearing aids and cochlear implants (*23-25*). The hypothesis that frogs’ lungs function in creating SCE stems from early qualitative reports that tympanic sensitivity can vary as a function of lung inflation (*16-18*). The SCE hypothesis predicts that sound transmission through inflated lungs to the middle ears has one or both of two effects on hearing in frogs. It could selectively augment the ability of sound frequencies emphasized in conspecific advertisement calls to drive vibrations of the tympanum. Alternatively, it could selectively reduce the tympanum’s sensitivity to non-call frequencies that may be characteristic of sources of environmental noise. We tested these two predictions using the American green treefrog (*Hyla cinerea*; Hylidae), a species in which males call to attract females in noisy breeding choruses, often with multiple other frog species, and females choose their mate based on perceived features of his calls.

## RESULTS

### Inflated lungs selectively reduce tympanum vibrations to non-call frequencies

We used laser vibrometry to assess the impacts of lung inflation on the tympanum’s sensitivity across the frequency spectrum and across the horizontal sound field. Using temporarily immobilized females as subjects, we compared the vibration amplitudes of the tympanum in response to free-field acoustic stimulation by broadcasting a frequency modulated (FM) sweep from each of 12 sound incidence angles surrounding the animal (Fig. S1A). Measurements were repeated with the lungs in a naturally inflated state and after manual deflation of the lungs (*n* = 21; Fig. 1C). The tympanum was most responsive to frequencies between about 1000 and 5000 Hz (Figs. 2A, 2B, S1A). However, in response to frequencies in the range of 1400 to 2200 Hz, tympanum vibration amplitude was substantially reduced, on average by 4 to 6 dB, when the lungs were inflated compared with the deflated state (Fig. 2C). Within this frequency range, the maximum reduction in tympanum vibration amplitude, averaged across individuals (mean ± 95% CI), was 10.0 ± 1.8 dB (range: 3.3 to 17.4 dB) and was significantly nonzero (two-tailed, one-sample t test: t_20_ = 10.91, p < 0.001). While the reduction in vibration amplitude spanned the frontal hemifield (Fig. 2C), its bandwidth and magnitude were larger when sound originated from within the contralateral portion of the frontal hemifield (i.e., between 0° and -90°). The modal and median sound incidence angles corresponding to the maximum reduction in tympanum vibration amplitude were -60° and -30°, respectively.

**Fig. 2.**
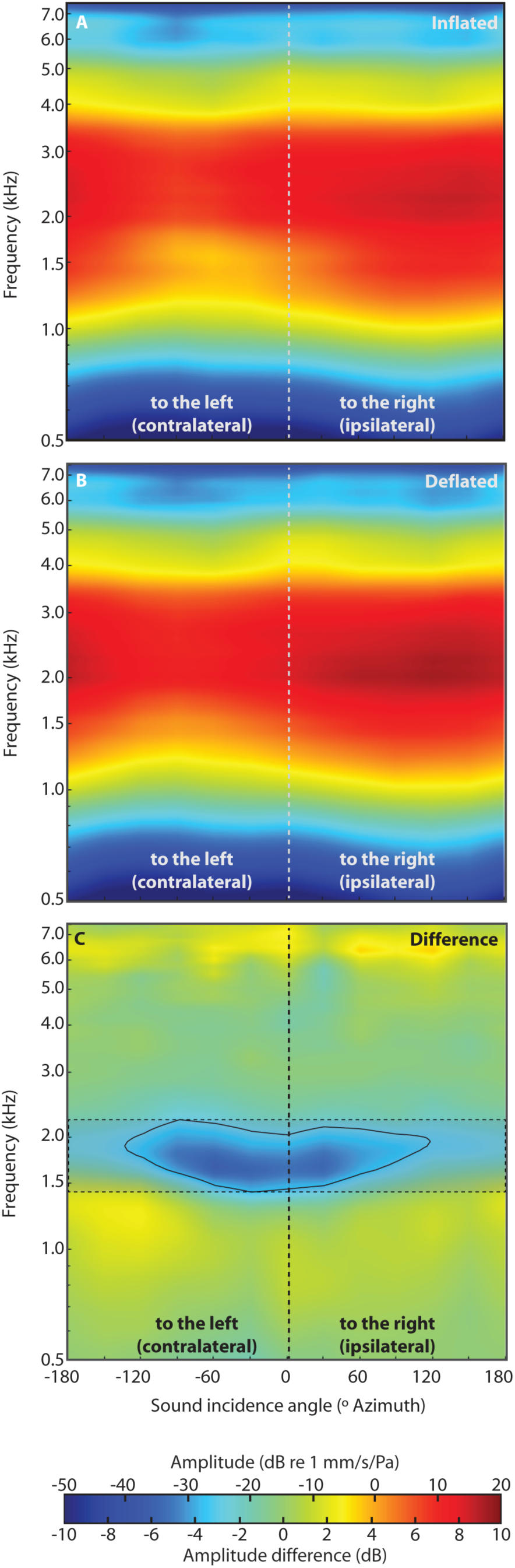
Inflated lungs selectively reduce tympanum vibrations in the frequency range of 1400 to 2200 Hz. **(A-B)** Heatmaps showing the mean vibration amplitudes of the right tympanum measured using laser vibrometry in response to free-field acoustic stimulation by an FM sweep presented from each of 12 sound incidence angles (0° to ±180° in 30° steps) surrounding the animal (Fig. S1A). Measurements were repeated in the inflated **(A)** and deflated **(B)** states of lung inflation (*n* = 21 individuals). **(C)** Heatmap showing the mean differences between the vibration amplitudes of the tympanum in the inflated and deflated states of lung inflation (inflated – deflated) across frequency and sound incidence angle (*n* = 21 individuals). The black contour encloses frequencies and angles where inflated lungs reduced vibration amplitudes by ≥ 4 dB. The minimum and maximum frequencies enclosed by the contour are 1400 Hz and 2200 Hz, respectively. The dashed lines and shaded box enclose frequencies between 1400 Hz and 2200 Hz across all angles and are reproduced in subsequent figures.

We found no evidence that inflated lungs selectively augment the ability of frequencies emphasized in conspecific advertisement calls to drive the tympanum. Similar to other frogs in the genus *Hyla* (*9*), male green treefrogs produce an advertisement call with a frequency spectrum consisting of two prominent spectral peaks that are analogous to the formant frequencies present in human vowel sounds (*26*). The low-frequency peak is important for long-distance communication (*27*) and source localization (*28, 29*), whereas the high-frequency peak appears to be more important in female mate choice (*27, 30*). In a sample of 457 advertisement calls (≅ 20 calls from each of 23 males), the mean (± SD) frequencies of the two peaks were 834 ± 14 Hz and 2730 ± 34 kHz (Fig. 3A). It is worth noting that the mean (± SD) frequency of the prominent valley in the frequency spectrum between these two spectral peaks was 1653 ± 39 Hz, which falls within the range of frequencies (1400 to 2200 Hz) where inflated lungs reduce tympanic sensitivity (Fig. 3A). At the two spectral peaks of conspecific calls, the tympanum’s vibration amplitude did not vary as a function of lung inflation. Across angles of sound incidence, the mean (± 95 CI) magnitude of the tympanum response at frequencies of 834 Hz and 2730 Hz differed between the inflated and deflated lung states by 0.7 ± 1.8 dB (Fig. 3B) and 0.8 ± 1.8 dB (Fig. 3C), respectively; these mean values were not significantly different from zero (two-tailed, one-sample t tests: t_20_ = 0.8 and t_20_ = 0.9, respectively, ps > 0.393). Overall, there was little evidence that inflated lungs augmented the tympanum’s response at any specific combinations of frequency and location. Tympanum responses were slightly higher (by 1.3 to 2.7 dB on average) at frequencies between 900 and 1500 Hz, but only in a restricted part of the contralateral sound field behind the animal, between about -90° and -180° (Fig. 2C). Other lung-mediated increases in tympanum sensitivity were similarly small and occurred at high frequencies (e.g., between 6000 and 7000 Hz; Fig. 2C) at which both the tympanum and the peripheral auditory nervous system are not very responsive (cf. Figs. 2A, 2B, S1A) (*31, 32*).

**Fig. 3.**
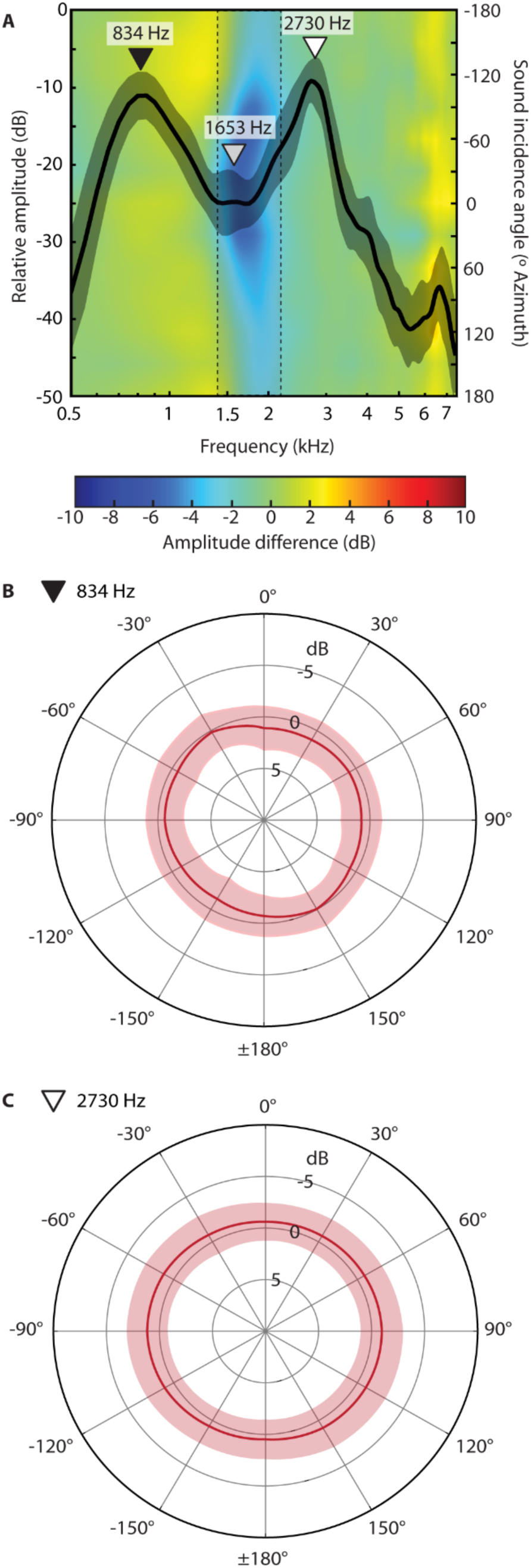
Inflated lungs selectively reduce tympanum vibrations to non-call frequencies. **(A)** Frequency spectrum of the green treefrog advertisement call overlaid on a heatmap showing lung-mediated effects on tympanic sensitivity (redrawn from Fig. 2C and rotated 90° clockwise). The spectrum consists of two prominent spectral components, a low-frequency peak (834 Hz, black triangle) and a high-frequency peak (2730 Hz, white triangle) separated by a prominent valley in the spectrum centered at 1653 Hz (gray triangle). The thick black line and shaded gray area depict the mean ± 1 SD call spectrum (*n* = 23 males, 457 individual calls). The dashed lines and shaded box enclose frequencies between 1400 Hz and 2200 Hz across all angles. (**B, C**) Polar plots showing the mean ± 95% CI difference (in dB) in the tympanum’s response between the inflated and deflated conditions (inflated – deflated) at the frequencies of the two spectral components of conspecific calls. The 95% CIs included 0 dB at all angles of sound incidence.

Together, these data suggest the inflated lungs of frogs function in creating a directionally tuned “notch filter” at the level of the tympana that attenuates a narrow range of sound frequencies predominantly within each contralateral frontal field. Because of this selectivity, the tympanum’s sensitivity to the spectral peaks of vocal signals was independent of the state of lung inflation. In fact, the spectrum of the advertisement call has a prominent valley that lacks sound energy at the very frequencies impacted by the lungs.

### Lung resonance generates a directionally tuned tympanic notch filter

Because the frog’s lungs are large air-filled cavities overlain by a relatively thin body wall, they resonate in response to free-field acoustic stimulation (*15, 19*). We hypothesized that the lungs’ resonance functions in generating the notch filter that selectively reduces the tympanum’s sensitivity. This hypothesis predicts that the peak resonance frequency of the lungs should correspond closely to the frequency range of reduced tympanum sensitivity observed when the lungs are inflated (1400 to 2200 Hz) and that the resonance of the lungs should have a subtractive effect on the tympanum’s response within this frequency range.

Biophysical measurements revealed that the lungs exhibit a prominent resonance that corresponds closely to the frequency range of the lung-mediated reduction in tympanum sensitivity. We used laser vibrometry to quantify the vibration amplitude of the body wall overlying the lungs in response to free-field acoustic stimulation by an FM sweep broadcast from an ipsilateral position (*n* = 10; Fig S1B). The measured lung resonance had a mean (± 95% CI) peak frequency of 1558 ± 89 Hz (range: 1400 Hz to 1850 Hz) and mean (± 95% CI) 10-dB down points of 1244 ± 76 Hz and 1906 ± 151 Hz (Fig. 4A). There was a significant negative correlation between peak resonance frequency and body size, as expected if lung volume varies directly with body size (Fig. S2; two-tailed Pearson r = -0.705, R^2^ = 0.498, p = 0.0228). After manually deflating the lungs, the mean (± 95% CI) magnitude of the peak resonance frequency was attenuated by 57.6 ± 6.0 dB and was significantly lower than the magnitude measured in the inflated condition (Fig. 4A; two-tailed, one-sample t test: t_9_ = 18.7, p < 0.0001). Following manual reinflation of the lungs, the mean (± 95% CI) peak frequency of the lung resonance was restored (1627 ± 98 Hz; range: 1450 Hz to 1928 Hz) and did not differ from that measured in the inflated state (Fig. 4A; two-tailed, paired t test: t_9_ = 1.88, p = 0.0922). These laser measurements, thus, confirmed that under free-field acoustic stimulation, the resonance frequency of inflated lungs corresponds closely to the frequency range of reduced tympanum vibrations when the lungs are inflated.

**Fig. 4.**
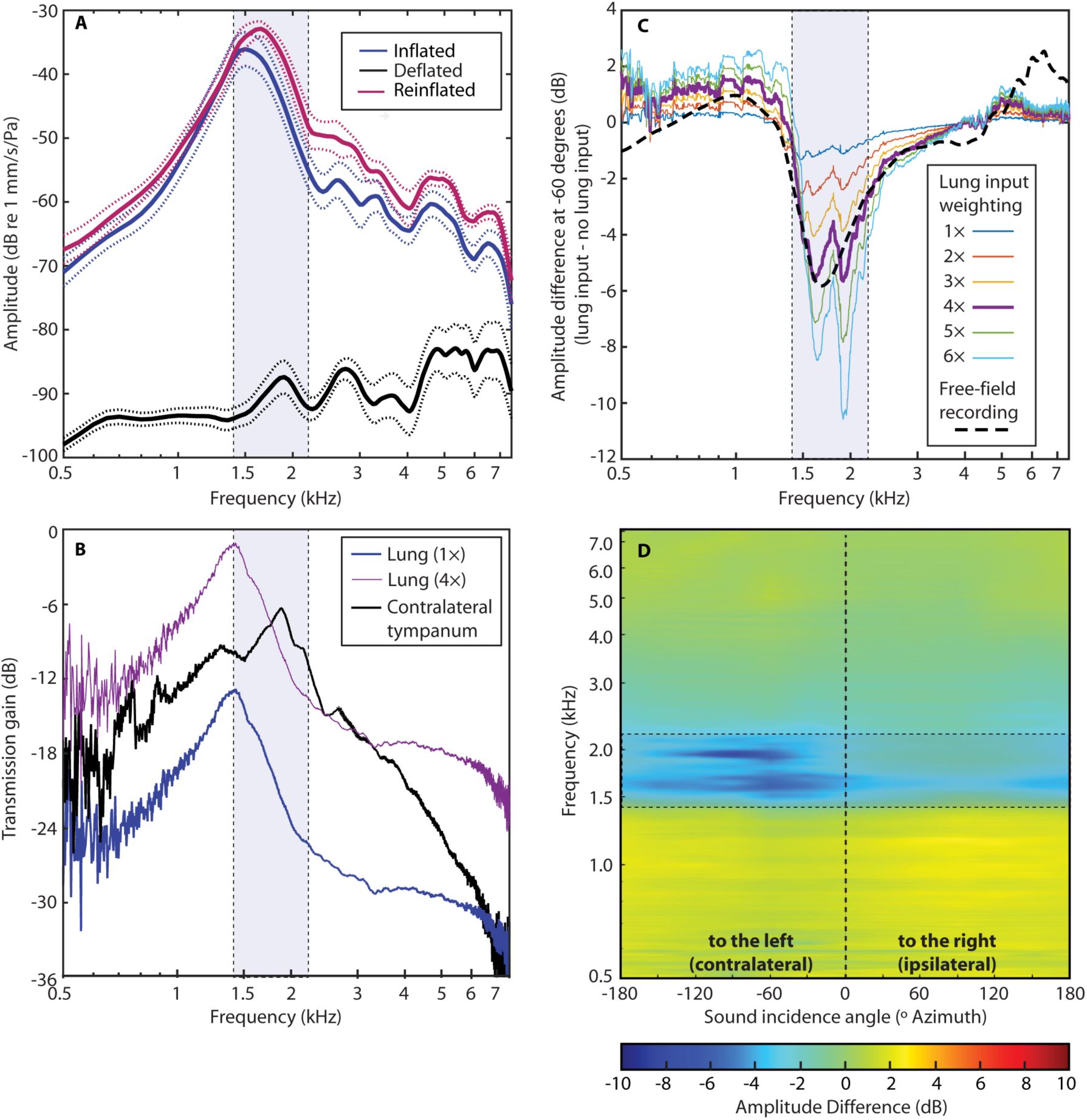
Lung resonance generates a tympanic notch filter. **(A)** Under free-field acoustic stimulation by an FM sweep, the inflated lungs of green treefrogs resonate at frequencies between 1200 to 1900 Hz. Depicted here are the mean (solid lines; *n* = 10 individuals) ± 95% CIs (dashed curves) vibration amplitudes of the body wall above the right lung in the inflated, deflated, and reinflated conditions. **(B)** Transmission gain of indirect sound input to the internal surface of the ipsilateral tympanum from the lungs (TG_L_, with 1× and 4× transmission gain weightings) and contralateral tympanum (TG_C_) measured under conditions of local acoustic stimulation by an FM sweep (Fig. S1C). **(C)** Predicted effects of inflated lungs based on reconstructing the tympanum’s free-field response at a contralateral sound incidence angle of - 60° using measures of transfer functions obtained under conditions of local acoustic stimulation by an FM sweep. Differences between reconstructed tympanum responses for the inflated and deflated conditions are shown for six transmission gain weightings (1× to 6×) of the lung input to account for the fact that the lungs would be stimulated to a much larger degree under free-field acoustic conditions. The actual impacts of the lung on the tympanum’s free-field response are shown by the dashed black line (redrawn from the -60° contour in the heatmap in Fig. 2C), which closely matches the predicted response for a lung transmission gain weighting of 4×. (**D**) Predicted effects of inflated lungs on the tympanum’s free-field response at all angles of sound incidence based on measures of transfer functions under conditions of local acoustic stimulation by an FM sweep and using a lung input weighting of 4×. In all panels **(A-D)**, the dashed lines and shaded blue rectangle enclose frequencies between 1400 Hz and 2200 Hz.

We tested the prediction that inflated lungs have a frequency-specific subtractive effect by comparing reconstructions of the tympanum’s free-field response with the lungs inflated versus deflated. To do so, we used local acoustic stimulation (Fig. S1C) to quantify the transmission gain (TG) of indirect sound input to the internal surface of the tympanum via the inflated lungs (TG_L_) and the contralateral tympanum (TG_C_) (*33*). Transmission gain represents a relative measure of how efficiently each indirect sound input is coupled to the tympanum. At TG = 0 dB, sounds impinge on the internal and external surfaces of the tympanum with equal amplitudes and interact depending on their relative phases; negative values indicate sound reaches the internal surface of the tympanum at a relatively lower amplitude. The median peak TG_L_ occurred at 1430 Hz (Fig 4B; *n* = 6 individuals), which falls within the range of the lung’s peak resonance frequencies measured with free-field acoustic stimulation (1400 Hz to 1850 Hz; cf. Fig. 4A). The peak TG_L_ was -13 dB (i.e., 13 dB lower) compared with the direct sound input to the tympanum’s external surface. The median peak TG_C_ occurred at 1852 Hz and was -6 dB (i.e., 6 dB lower) compared to direct sound input to the tympanum’s external surface (Fig. 4B).

Reconstructed tympanic responses to free-field stimulation exhibited the predicted lung-mediated reduction in vibration amplitude in the range of 1400 to 2200 Hz (Fig. 4C). Responses in the deflated state were reconstructed after excluding the lung input from all computations. To reconstruct responses with inflated lungs, we explored a range of transmission gain weightings for TG_L_ because local stimulation of the body wall (Fig. S1C) underestimates its real magnitude due to the much larger surface area of the body wall exposed to sound during free-field stimulation. At a TG_L_ weighting of 4×, tympanum responses reconstructed for the inflated lung condition were reduced, relative to the deflated condition, by approximately 4 to 6 dB at -60° in the frequency range of 1400 to 2200 Hz. This reduction is similar in magnitude to that observed in measured free-field responses (Fig. 4C). When this lung-mediated effect was modeled across additional angles of sound incidence, differences in the tympanum’s reconstructed free-field responses between the inflated and deflated states revealed broad patterns consistent with those determined with free-field acoustic stimulation: tympanum sensitivity was reduced within the frequency range of 1400 to 2200 Hz when the lungs were inflated, and these reductions were most pronounced at contralateral angles (cf. Figs. 2C and 4D). A transmission gain weighting of 4× for TG_L_ corresponds to a lung transmission gain amplitude that is 12 dB higher than the unweighted value of -13 dB reported above. Our reconstructions, therefore, suggest the peak transmission gain from the lungs is probably closer to -1 dB relative to the direct sound input to the tympanum’s external surface and +5 dB relative to the peak transmission gain amplitude from the contralateral tympanum (Fig. 4B). Hence, sound transmission to the internal surface of the tympanum through the lungs is quite substantial and, at the lung resonance frequency, it is nearly equivalent in magnitude to direct sound stimulation of the tympanum’s external surface.

Together, bioacoustic and biophysical measurements, combined with reconstructing free-field responses, provide direct evidence that the lung resonance generates a directionally tuned “notch filter” that attenuates the contralateral portion of each tympanum’s frontal field response within a narrow range of frequencies between 1400 to 2200 Hz. But how might lung mediated notch filtering function in hearing and sound communication? To answer this question, we examined the lungs’ impacts on tympanic responses in relation both to a physiological model of peripheral frequency tuning in frogs and to the frequency spectra of common sources of environmental noise.

### Lung mediated notch filtering sharpens peripheral frequency tuning

We modeled peripheral frequency tuning in green treefrogs by estimating frequency tuning curves as 4th order gammatone filters that we parameterized using published data on best excitatory frequencies, bandwidths, and thresholds for 172 auditory nerve fibers in this species (*31*) (Fig. 5). Separate populations of auditory nerve fibers innervating the amphibian papilla are tuned to low frequencies (up to about 700 Hz) and mid-range frequencies (up to about 1300 Hz), whereas a third population of auditory nerve fibers innervating the basilar papilla is more consistently tuned to a common range of higher frequencies (Fig. 5). Consistent with the notion of matched filtering in the spectral domain, the thresholds of auditory nerve fibers innervating the amphibian and basilar papillae are lowest at frequencies matching, respectively, the low-frequency and high-frequency spectral peaks in the advertisement call (Fig. 5).

**Fig. 5.**
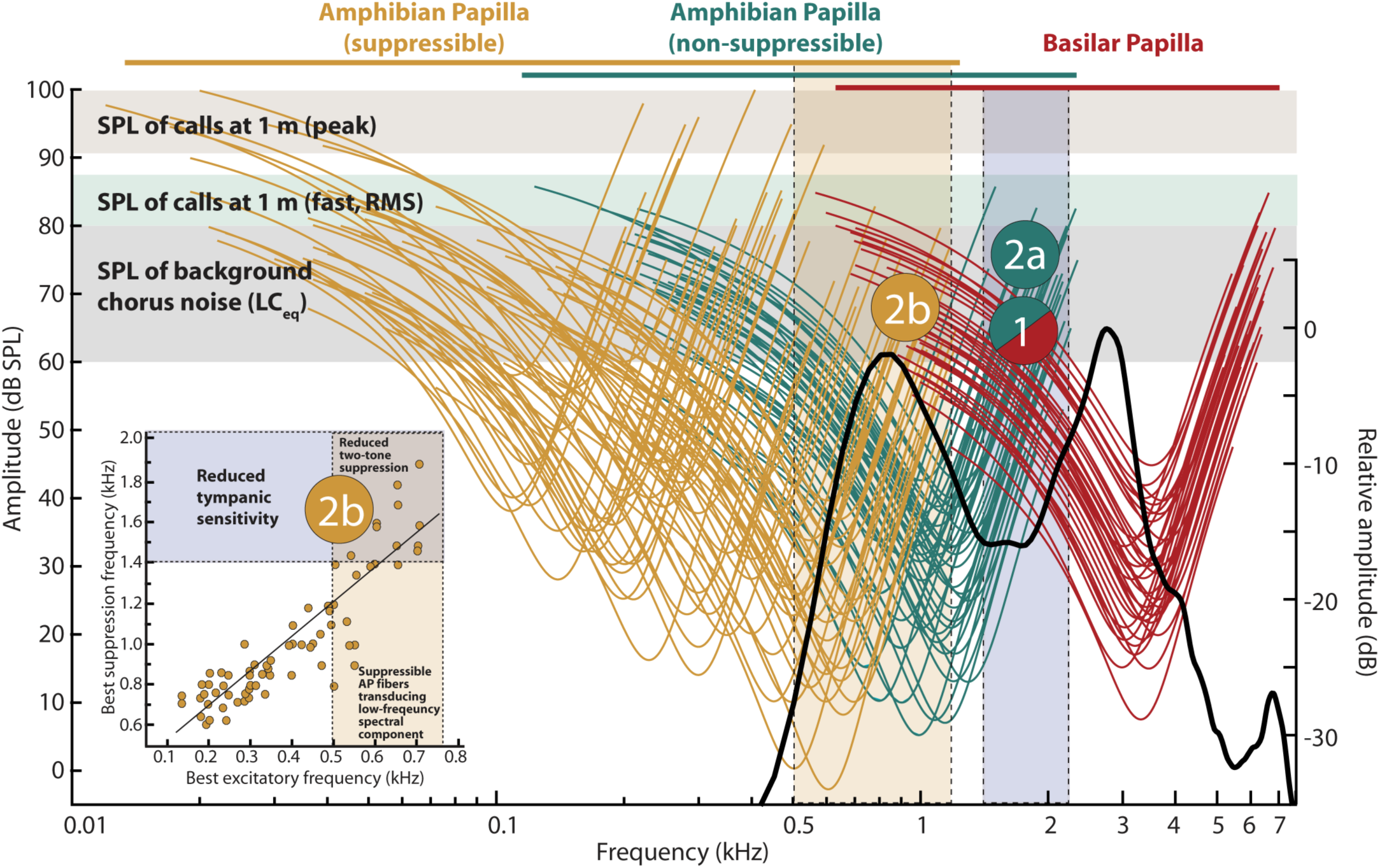
Lung mediated notch filtering sharpens peripheral frequency tuning and reduces energetic masking and two-tone rate suppression. Modeled tuning curves for 172 auditory nerve fibers in green treefrogs are shown in relation to the frequency range of lung-mediated reductions in tympanum sensitivity (1400 to 2200 Hz, right shaded blue rectangle), the spectrum of conspecific calls (solid black line redrawn from Fig. 3A), and the sound pressure levels (SPLs) of conspecific advertisement calls (*49*) and background chorus noise for a closely-related treefrog (*50*). Tuning curves are depicted separately for low-frequency and mid-frequency fibers innervating the amphibian papilla (AP) and for high-frequency fibers innervating the basilar papilla (BP). Neural responses of low-frequency fibers innervating the AP can be suppressed by frequencies in the range of mid-frequency AP fibers (*36*). The inset shows the best suppression frequency for a given best excitatory frequency for 70 auditory nerve fibers derived from the AP having best excitatory frequencies at or below approximately 700 Hz (redrawn from (*36*)). The greatest two-tone suppression of excitatory responses to a tone presented 10 dB above threshold at a unit’s best excitatory frequency was observed when a second tone was simultaneously presented at the unit’s best suppression frequency. As the modeled tuning curves illustrate, nerve fibers with best excitatory frequencies in the range of 0.5 to 0.7 kHz transduce the low-frequency spectral component of conspecific calls at the high sound amplitudes used for communication (left shaded gold rectangle). Approximately half of the nerve fibers with best excitatory frequencies within this range can be suppressed by sound energy in the range of 1400 Hz to 1900 Hz. The model predicts two mechanisms by which a reduced tympanum response to frequencies between 1400 Hz and 2200 Hz is expected to improve sensory processing of conspecific calls. Reduced stimulation by environmental noise of both non-suppressible AP fibers and BP fibers at frequencies where their tuning overlaps (‘1’) would reduce energetic masking of both spectral components in conspecific calls. Reduced stimulation of non-suppressible AP fibers (‘2a’) would additionally reduce two-tone rate suppression of high-frequency, suppressible AP fibers (‘2b’) that transduce the low-frequency component of the call.

The key finding revealed by this physiological model is that lung-mediated impacts on tympanic sensitivity occur in the frequency range where the tuning of the amphibian and basilar papillae can overlap (Fig. 5). At low amplitudes near threshold, neither inner ear papilla responds to frequencies between 1400 and 2200 Hz. However, the bandwidth of individual auditory nerve fibers broadens considerably at higher sound levels more typical of communication in natural environments, ultimately causing the tuning of the two inner ear papillae to overlap (*32, 34*). Much of the frequency range of overlap between the amphibian and basilar papillae at high sound levels corresponds to the frequency range (1400 to 2200 Hz) where notch filtering by inflated lungs reduces the tympanum’s vibration amplitude (Fig. 5). Given this correspondence, the modeled tuning of auditory nerve fibers reveals two possible mechanisms by which the subtractive notch filtering generated by inflated lungs could function to counteract the negative impacts of environmental noise in the frequency range of 1400 to 2200 Hz.

First, inflated lungs would reduce energetic masking of conspecific advertisement calls. Consistent with behavioral studies (*34*) and recordings of auditory brainstem responses (*32*), auditory nerve fibers innervating both the mid-frequency region of the amphibian papilla and the basilar papilla are predicted to respond to frequencies in the range of 1400 to 2200 Hz at high amplitudes typical of communication. By selectively reducing the sensitivity of the tympanum, notch filtering by inflated lungs would reduce the ability of environmental noise in this frequency range to drive auditory nerve responses that could otherwise mask neural responses to conspecific calls. Second, notch filtering by inflated lungs would reduce two-tone rate suppression along the amphibian papilla. Two-tone rate suppression is a well-known feature of auditory processing in vertebrates (*35*), including green treefrogs (*36, 37*), whereby neural responses to one frequency are suppressed by the addition of a second frequency. In green treefrogs, suppressible auditory nerve fibers with best frequencies between approximately 500 and 700 Hz respond to the low-frequency component of conspecific calls at the high amplitudes typical of communication (Fig. 5). Importantly, these fibers are suppressed by simultaneous sound energy between approximately 1400 and 2000 Hz (see Fig. 5 *inset*) (*36, 37*). Thus, notch filtering that selectively attenuates the tympanum’s response in this frequency range should additionally reduce two-tone rate suppression of excitatory neural responses to conspecific calls.

Our physiological model suggests lung mediated notch filtering functions generally to sharpen peripheral frequency tuning in the range of 1400 to 2200 Hz. This interpretation is consistent with the SCE hypothesis: by selectively attenuating the tympanum’s response to frequencies in the valley between the peaks in the conspecific call spectrum, inflated lungs improve the “match” of the peripheral matched filter to frequencies in conspecific signals. This improved match should both reduce energetic masking and two-tone rate suppression that could otherwise impair neural processing of the spectral peaks of conspecific advertisement calls, thereby increasing the neuronal signal-to-noise ratio for communication. A wider band of notch filtering would interfere with tympanic transmission of the spectral peaks in conspecific calls. An important remaining question, therefore, is what sources of environmental noise in the range of 1400 to 2200 Hz would be attenuated by lung mediated notch filtering?

### Lung mediated notch filtering improves the signal-to-noise ratio for communication

For frogs that call in multi-species choruses, the calls of other species breeding at the same times and places represent prominent and behaviorally relevant sources of environmental noise that can interfere with communication (*5, 9*). Therefore, we evaluated the extent to which the calls of syntopically breeding heterospecific frogs constitute environmental noise in the frequency range attenuated by the lungs. To this end, we integrated analyses of continent-scale citizen science data from the North American Amphibian Monitoring Program (NAAMP) (*38*) with bioacoustic analyses of archived recordings of frog calls. NAAMP was a long-term (1994-2015) effort to monitor frog populations using roadside calling surveys conducted across 26 states that encompass most of the green treefrog geographic range in the eastern, central, and southern United States. In the NAAMP dataset, there were 19,809 reports of “co-calling” between green treefrogs and a total of 42 other species, meaning these 42 species were observed to call at the same times and places as our focal species (Fig. S3).

Social network analysis revealed that just 10 of the 42 co-calling heterospecific species (i.e., 24% of all co-calling species) accounted for the overwhelming majority (79%) of the 19,809 observed instances of co-calling between green treefrogs and one or more other species in the NAAMP dataset (Figs. 6 and S3). Of these top-10 heterospecific species, half (i.e., just 5 of 42 total heterospecific species) accounted for 42% of all instances of co-calling and produce advertisement calls with prominent spectral peaks falling in the range of 1400 to 2200 Hz (Fig. 6). One of these five species, for example, is Fowler’s toad (*Anaxyrus fowleri*), which produces a long, loud call with a single spectral peak in the range of 1400 to 2200 Hz (Fig. 6). Others of these five species, such as the barking treefrog (*Hyla gratiosa*), the North American bullfrog (*Lithobates catebeianus*), and the green frog (*Lithobates clamitans*) produce calls with bimodal spectra in which one of two spectral components falls within the range of 1400 to 2200 Hz (Fig. 6). We selected these five species’ calls for further investigation.

**Fig. 6.**
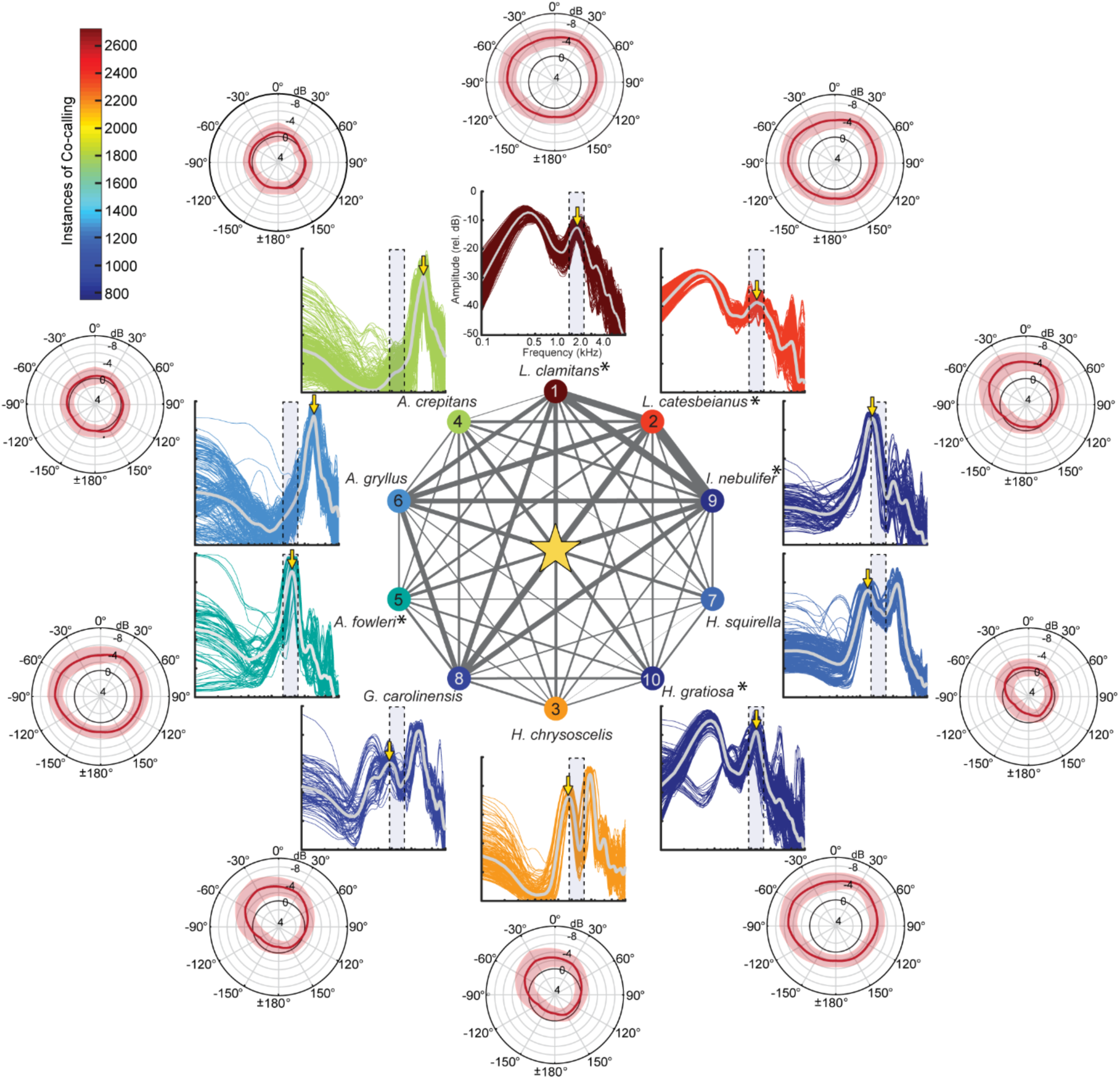
Lung mediated notch filtering improves the signal-to-noise ratio for communication by reducing the tympanum’s response to environmental noise. The central figure depicts the social network of co-calling between green treefrogs (star) and the top-10 heterospecific species identified from the NAAMP dataset (Fig. S3). Network edges reflect the relative number of observations of co-calling. Five of the top-10 species produce calls with substantial acoustic energy in the range of reduced tympanum vibration amplitude (asterisks; Table S1). Surrounding the network are depictions of the frequency spectra of each heterospecific species’ advertisement calls. Gray lines depict the mean spectrum, averaged over all individuals and calls; colored lines depict the spectrum for each analyzed call and are scaled in color according to the instances of co-calling with green treefrogs (color bar). The shaded blue rectangle in each spectrum represents the frequency range of maximal reduction in the green treefrog’s tympanum vibration amplitude (1400 to 2200 Hz) as a result of lung inflation. Surrounding the spectra are polar plots depicting the mean ± 95% CI difference (in dB) in the tympanum’s response between the inflated and deflated conditions (inflated – deflated) at the frequency indicated by a downward pointing yellow arrow on each spectrum (*n* = 21 individuals).

To model the impacts of lung mediated notch filtering on the tympanum’s response to these five species’ calls, we passed the spectra of their advertisement calls through simulated tympanic filters corresponding to 0° (frontal) and -30° and -60° (contralateral) with the lungs inflated versus deflated. We then computed the difference in the tympanum’s predicted vibration amplitude between states of lung inflation at the spectral peak of each heterospecific species’ calls within or closest to the range of 1400 to 2200 Hz. Consistent with a functional role in noise reduction, the notch filtering generated by inflated lungs reduced the tympanum vibration amplitude in response to the relevant spectral peaks of these five species’ calls by approximately 4 to 6 dB (Fig. 6; Table S1). The greatest lung-mediated reductions in tympanic responses in the range of 1400 to 2200 Hz occurred for the calls of green frogs (*L. clamitans*), bullfrogs (*L. catesbeianus*), and barking treefrogs (*H. gratiosa*) (Fig. 6; Table S1). Notably, bullfrogs and green frogs co-called with green treefrogs most frequently in the NAAMP dataset, with these two heterospecific species accounting for 26% of all reported instances of co-calling. Barking treefrogs are the sister species of green treefrogs. The costs of hybrid matings with barking treefrogs have driven evolutionary change in the spectral preferences of female green treefrogs (*39*); therefore, mitigating the risk of mis-mating could be one additional benefit of the lungs’ impacts on tympanic responses. Taken together, these data are consistent with the interpretation that SCE created by lung mediated notch filtering improves the signal-to-noise ratio for communication by selectively reducing the tympanum’s sensitivity to prominent sources of environmental noise that include the calls of heterospecific frog species in multi-species choruses.

## DISCUSSION

Resonances of air-filled structures such as lungs or swim bladders improve sound detection in aquatic vertebrates (*40*), and sound detection via the lungs also played important roles in hearing during the evolutionary transition of vertebrates from water to land (*20*). Thus, it is likely that the lungs of the earliest terrestrial vertebrates functioned as accessory sound receiving structures prior to the subsequent evolution of tympanic middle ears (*12*) and acoustic communication (*11*). Our data suggest evolution has co-opted this ancient adaptation for sound reception through the lungs to provide spectral contrast enhancement that improves the signal-to-noise ratio for communication in modern frogs. The inflated lungs of female green treefrogs act as resonators that reduce the tympanum’s sensitivity to sounds occurring within a narrow but biologically important range of frequencies. This frequency range, which falls precisely in the valley between the two spectral peaks of the species-specific advertisement call, encompasses frequencies that are transduced by both inner ear papillae and that are used by some of the most frequently encountered heterospecifics in multi-species frog choruses. In essence, inflated lungs enhance spectral contrast by sharpening the peripheral matched filter. In turn, inflated lungs should improve the signal-to-noise ratio for vocal communication by reducing energetic masking and two-tone rate suppression by sources of environmental noise that potentially interfere with processing conspecific calls. Thus, frogs’ lungs help them solve a multi-species cocktail party problem.

This study’s support of the SCE hypothesis suggests the lungs in one major clade of tetrapods serve a heretofore unknown noise-control function in vertebrate hearing and sound communication. Moreover, the impact of the frog’s inflated lungs on its tympanum vibrations bears striking similarity to SCE in the field of human hearing and speech communication (*23-25*). People with sensorineural hearing loss have difficulties understanding speech in noisy social settings that stem, in part, from having broader auditory filters (i.e., reduced frequency selectivity). SCE algorithms can produce a 2 to 4 dB increase in the signal-to-noise ratio that yields significant improvements in word recognition and response times in hearing impaired listeners (*24*). Some SCE algorithms that improve speech recognition for cochlear implant users do so by attenuating the valleys between the peaks of formant frequencies in the speech spectrum without altering the levels of the peaks themselves (*23*). This human engineering solution is functionally similar to how inflated lungs impact tympanic sensitivity in frogs: the state of lung inflation had no impact on the formant-like spectral peaks present in conspecific calls, but inflated lungs attenuated (by up to 10 dB on average) the tympanum’s response to frequencies in the valley between them. Notably, lung-mediated notch filtering occurred in the frequency region (1400 to 2200 Hz) where, at high sound levels, the spectral tuning of the separate inner ear organs is broader and frequency selectivity is reduced. In a closely related treefrog, even large changes in lung volume (e.g., 66.7% reduction) yielded no change in the magnitude of the lung’s resonance and produced only a small change (e.g., 100 Hz increase) in the peak resonance frequency (*16*). These findings suggest the acoustical properties of the lung input probably change very little over the respiratory cycle. Hence, the lung-to-ear transmission pathway in frogs provides a stable biophysical mechanism for real-time SCE that begins at the tympanum itself. In green treefrogs, such a mechanism should have the effect of enhancing call recognition, though testing this mechanism directly may be impossible due to the difficulty of manipulating lung volume in behaving animals engaged in call recognition tasks. Previous work, however, lends indirect support to the operation of such a mechanism, as call recognition is enhanced when both spectral peaks are present and degraded when additional spectral peaks in the range of 1400 to 2200 Hz are artificially added to calls (*26, 37*).

A mechanism for SCE might be particularly beneficial to frogs by mitigating negative impacts of the environmental noise created by the calls of other species in a multi-species chorus. Because accurate perception of sexual signals is often tightly linked to evolutionary fitness, species that communicate acoustically in multi-species breeding aggregations can be under intense selection to solve cocktail-party-like problems (*1, 2, 4-7*). The negative impacts of auditory masking caused by background noise and overlapping signals should, thus, favor the evolution of adaptive mechanisms that promote more efficient communication in noise. Results from this study suggest the frog’s lung-to-ear sound transmission pathway functions as a novel receiver adaptation to mitigate problems of auditory masking that might be particularly detrimental to effective communication in multi-species choruses.

Our results also resolve a longstanding paradox in research on comparative vertebrate hearing. Previous studies of frogs (*13, 16-18*), including green treefrogs (*22*), indicate that the lung-to-ear sound transmission pathway sharpens the tympanum’s inherent directional tuning at frequencies corresponding to the lung resonance frequency, but that it has very little (if any) impact on directionality at the signal frequencies used for social and sexual communication with conspecifics. That the frog’s lungs improve directional hearing at frequencies not used for communication has remained paradoxical, because increased directionality at conspecific call frequencies would presumably confer selective advantages in the dark and physically complex environment of a nighttime chorus by improving a receiver’s ability to locate sources of conspecific calls (e.g., potential mates). A pattern observed but not considered in earlier studies (*16-18*) is that lung-mediated improvements in directionality occur because the tympanum’s sensitivity was reduced to sound frequencies near the lung’s peak resonance frequency in a direction dependent manner. Our findings resolve the apparent paradox by emphasizing that it is the frequency-dependent, lung-mediated decrease in tympanic sensitivity – and not the associated increase in directionality – that plays a functional role in hearing and sound communication.

In conclusion, we propose that natural selection has acted, at least in some frog species, to exploit the lung-to-ear coupling to facilitate perception of conspecific calls in noisy environments by providing a mechanism for real-time spectral contrast enhancement in the auditory periphery. That the frog’s lungs play a role in spectral contrast enhancement significantly broadens our understanding of the diversity of evolutionary adaptations to noise problems in nonhuman animals as well as the function of a unique sound transmission pathway in one of the most vocal groups of extant vertebrates.

## MATERIALS AND METHODS

### Animals

Subjects were 25 female green treefrogs collected on the grounds of the East Texas Conservation Center in Jasper County, Texas, U.S.A. (30°56’46.15”N, 94°7’51.46”W). Animals were housed in the laboratory on a 12-hour photoperiod, provided with access to perches and refugia, fed a diet of vitamin-dusted crickets, and given ad libitum access to fresh water. All procedures were approved by the Institutional Animal Care and Use Committees of the University of Minnesota (#1401-31258A) and complied with the NIH *Guide for the Care and Use of Laboratory Animals* (8^th^ Edition).

### Laser vibrometry measurements

We took laser measurements of 25 frogs (SVL: mean = 54.0 mm, range = 47.7 to 59.2 mm; mass: mean = 12.5 g, range = 8.3 to 17.6 g). For laser measurements, subjects were immobilized with succinylcholine chloride (5 μg/g). Over the 5-10 minutes during which the immobilizing agent took effect, subjects were allowed to regulate their own lung volume. After full immobilization was achieved, lung ventilation had stopped and lung inflation (based on body wall extension) resembled that observed for unmanipulated frogs sitting in a natural posture (*41*). We refer to this state of lung inflation as “inflated.” For some procedures, we also examined animals in one or two additional states of lung inflation that involved manually deflating and reinflating the lungs. To create a “deflated” condition, we expressed the air in the animal’s lungs by gently depressing the lateral body wall while holding the glottis open with the narrow end of a small, plastic pipette tip. We created a “reinflated” condition by gently blowing air by mouth through a pipette with its tip located just above the closed glottis; the movement of air was sufficient to open the glottis and inflate the lungs. We made every attempt to return the lungs to the natural level of inflation observed prior to manual deflation. When individuals were measured in multiple conditions, we always made measurements in the inflated condition first followed by the deflated condition and then the reinflated condition. While animals were immobilized, we periodically applied water to the dorsum to keep the skin moist to facilitate cutaneous respiration. All laser measurements of an individual subject were made during a single recording session of less than two hours. (Note that temporary manipulations of lung ventilation are possible in immobilized frogs because amphibians are capable of cutaneous gas exchange.) We excluded four animals from further analyses (final *n* = 21) because we could not visually confirm their state of lung inflation across treatments.

We conducted our experiments in a custom-built, semi-anechoic sound chamber with inside dimensions (L × W × H) of 2.9 m × 2.7 m × 1.9 m (Industrial Acoustics Company, North Aurora, IL). To reduce reverberations inside the chamber, the walls and ceiling were lined with Sonex acoustic foam panels (Model VLW-60; Pinta Acoustic, Inc. Minneapolis, MN). The floor of the chamber was covered in low-pile carpet. During recordings, subjects were positioned on a 30-cm tall pedestal made from wire mesh (0.9-mm diameter wire, 10.0-mm grid spacing). The tip of the subject’s mandible rested on a raised arch of thin wire, so that the animal sat in a typical posture in the horizontal plane with its jaw parallel to the ground and its head raised and in line with its body. The bottom of the pedestal was suspended 90 cm above the floor of the chamber using a horizontal, 70-cm long piece of Unistrut^®^ attached to its base (Unistrut, Harvey, IL). The Unistrut beam was mounted to a vibration isolation table (Technical Manufacturing Corporation, Peabody, MA) located against an inside wall of the chamber. The Unistrut and vibration isolation table were covered with the same acoustic foam that lined the walls and ceiling of the chamber.

We measured the vibration amplitude of the animal’s right tympanum or body wall using a laser vibrometer (PDV-100, Polytech, Irvine, CA). The laser was mounted on the same vibration isolation table from which the subject pedestal was mounted. We positioned the laser at approximately 70° to the animal’s right relative to the direction in which its snout pointed, which we consider to be 0° (Fig. S1). To enhance reflectance of the laser, a small (45 – 63 μm diameter), retroreflective glass bead (P-RETRO-500, Polytech, Irvine, CA) was placed at the center of the right tympanum and a position on the right, lateral body wall overlying the lung. The analogue output of the laser was acquired (44.1 kHz, 16 bit) using an external digital and analogue data acquisition (DAQ) device (NI USB 6259, National Instruments, Austin, TX) that was controlled by custom software written in MATLAB (v.2014a, MathWorks, Natick, MA) and running on an OptiPlex 745 PC (Dell, Round Rock, TX). The spectra of the acquired laser signals were calculated in MATLAB using the pwelch function (window size = 256, overlap = 50%). Laser spectra were corrected for small directional variation in the sound spectrum by subtracting (in dB) the spectrum recorded from a probe microphone from that of the acquired laser signal. In generating heatmaps of tympanum responses (e.g., Fig. 2), we used linear interpolation to determine vibration amplitude values at angles of sound incidence between those measured. Data between measurement points in polar plots were interpolated using a cubic spline.

### Free-field acoustic stimulation

Acoustic stimuli (44.1 kHz, 16-bit) were broadcast using the same software and hardware used to acquire the laser signal. We controlled signal levels for calibration and playback using a programmable attenuator (PA5, Tucker-Davis Technologies, Alachua, FL). The stimulus consisted of a frequency-modulated (FM) sweep that was 195 ms in duration, had linear onset and offset ramps of 10 ms, and linearly increased in frequency from 0.2 to 7.5 kHz over the 175-ms steady-state portion of its amplitude envelope. Responses were averaged over 20 repetitions of the stimulus. During measurements with the laser, we also recorded acoustic stimuli by positioning the tip of a probe tube of a G.R.A.S. 40SC probe microphone (G.R.A.S. Sound & Vibration A/S, Holte, Denmark) approximately 2 mm from the position on the animal’s body (e.g., the right tympanum or its right body wall) from which the laser recorded the response. The microphone’s output was amplified using an MP-1 microphone pre-amplifier (Sound Devices, Reedsburg, WI) and recorded using the NIDAQ device.

For free-field acoustic stimulation (Fig. S1A, S1B), the stimulus was amplified (Crown XLS1000, Elkhart, IN) and broadcast through a speaker (Mod1, Orb Audio, New York, NY) located 50 cm away from the approximate center of a subject’s head (measured along the interaural axis) as it sat on the pedestal. The speaker was attached to a rotating arm covered in acoustic phone and suspended from the ceiling of the sound chamber so that the center of the speaker was at the same height above the chamber floor as the subject and could be placed at any azimuthal position. We presented the stimulus from 12 different angles around the animal (0° to 330° in 30° steps; Fig. S1A). An angle of 0° corresponded to the direction in which the animal’s snout pointed, +90 corresponded to the animal’s right side (ipsilateral to the laser), and -90° corresponded to the animal’s left side (contralateral to the laser). For a given subject, we first recorded responses from the tympanum in the inflated condition. We began recordings at a randomly determined location around the subject and then recorded responses at each successive angle after repositioning the speaker in a counterclockwise direction. After making a recording from the 12th and final speaker location, we deflated the lungs and remeasured the tympanum beginning at the same randomly determined starting location used in the naturally inflated condition. For a subset of 10 subjects (SVL: mean = 54.4 mm, range = 47.7 to 59.2 mm; mass: mean = 13.1 g, range = 8.3 to 17.6 g), we also measured the vibration amplitude of the body wall overlying the lungs in the inflated and deflated states, as well as after manually reinflating the lungs, with the speaker positioned at +90°. We calibrated the FM sweep to be 85 dB SPL (sound pressure level re 20 µPa, fast, C-weighted) for each speaker position using a Brüel & Kjær Type 2250 sound level meter (Brüel & Kjær Sound & Vibration Measurement A/S, Nærum, Denmark) and a Brüel & Kjær Type 4189 ½-inch condenser microphone. For calibration, we suspended the microphone from the ceiling of the sound chamber by an extension cable (AO-0414-D-100) so that it hung at the position a subject’s head occupied during recordings.

### Transmission gain measurements and free-field reconstructions

We used laser vibrometry and local acoustic stimulation of the tympana and body wall to measure the transmission gain of sound input to the tympanum’s internal surface via the internally coupled tympanum and the lungs. For local acoustic stimulation (Fig. S1C), we broadcast the same FM sweep used in free-field stimulations through a Sony MDREX15LP earbud (Sony Corporation of America, New York, NY) positioned within approximately 2 mm of either the right tympanum (ipsilateral to the laser), the left tympanum (contralateral to the laser), or the left body wall overlying the left lung (contralateral to the laser) (Fig. S1C). The opening of the earbud was approximately the same diameter as the tympanum. The earbud was positioned using micromanipulators attached to a ring stand placed on the floor of the sound chamber. The tip of a metal probe tube connected to the G.R.A.S. 40SC probe microphone was inserted through the hybrid silicone of the earbud such that its opening barely protruded into the space between the speaker and the acoustically stimulated tympanum or body wall. The output of the probe microphone was amplified with the MP-1 microphone preamplifier and recorded with the NIDAQ data acquisition device. At each location of stimulation, we determined the mean vibration amplitude averaged over response to 20 repetitions of the stimulus with the lungs in the inflated condition.

Transmission gains were computed as follows. For each location of stimulation (Fig. S1C), we first computed a transfer function by dividing the average vibration spectrum of the right (ipsilateral) tympanum’s response recorded with the laser by the average sound spectrum at each location recorded with the probe microphone. Thus, separate transfer functions were computed for stimulation of the right tympanum (ipsilateral to the laser; H_I_(ω)), the left tympanum (contralateral to the laser; H_C_(ω)), and the body wall above the lung (contralateral to the laser; H_L_(ω)). From these transfer functions, we then computed the transmission gain for sound input to the internal surface of the ipsilateral tympanum via the contralateral tympanum as TG_C_ = H_C_(ω) / H_I_(ω) and via the lung-to-ear pathway as TG_L_ = H_L_(ω) / H_I_(ω).

Transfer functions measured with local acoustic stimulation allowed us to reconstruct the tympanum’s free-field response to sound. To accomplish this, we added the sounds arriving at the tympanum’s internal surface via the internally coupled, contralateral tympanum and via the lung-to-ear pathway to the sound measured at the external surface the tympanum. The sound arriving via the contralateral tympanum was computed as the sound impinging on the external surface of the contralateral tympanum (measured with the probe microphone) multiplied by the transmission gain of the contralateral tympanum (TG_C_). The sound arriving via the lung-to-ear pathway was computed as the sound impinging on the body wall (measured with the probe microphone) multiplied by the transmission gain of the lung-to-ear pathway as (TG_L_). Sound spectra were converted to complex numbers such that all addition was done vectorially and, thus, the resulting sum depended on both the amplitude and phase of each frequency. This sum of sound inputs was then multiplied by the transfer function of the ipsilateral tympanum (H_I_(ω)) to arrive at the predicted free-field transfer function. A range of weightings for TG_L_ (1× to 6×) were explored in reconstructing free-field responses because local acoustic stimulation underestimates the real magnitude of TG_L_ due to the smaller surface area of the body wall that is stimulated compared with free-field acoustic stimulation.

### Bioacoustic analyses

We made acoustic recordings of green treefrogs in their natural habitat during active breeding choruses. Between 15 May and 3 July 2013, we recorded 457 advertisement calls from 23 males that were calling in ponds at the East Texas Conservation Center. These recordings (44.1 kHz, 16 bit) were made between 2300 and 0200 hours using a Marantz PMD670 recorder (Marantz America, LLC., Mahwah, NJ) and handheld Sennheiser ME66/K6 microphone (Sennheiser USA, Old Lyme CT) held approximately 1 m from the focal animal. Male green treefrogs had a mean (± SD) snout-to-vent length (SVL) of 51.6 ± 3.9 mm and were recorded at a mean air temperature of 20.3 ± 3.0 °C. Recordings of green frogs (*Lithobates clamitans*) (*42*) and bullfrogs (*Lithobates catesbeianus*) (*43*) were obtained during previous studies by one of the authors (MAB); the calls of the remaining eight frog species included in this study were obtained from the Macaulay Library at the Cornell Lab of Ornithology. For each species, we analyzed 6 to 23 calls per male (median = 20 calls per male) for each of 9 to 25 males (median = 20 males per species) by computing the power spectrum of each call using MATLAB’s pwelch function (window size = 256, overlap = 50%). We determined the average call spectrum for a species by first averaging over the calls recorded from each individual and then over all individuals. Only recordings that were of sufficiently high signal-to-noise ratio and free from excessive background noise were included in these analyses.

### Model of peripheral frequency selectivity

We modeled frequency selectivity in green treefrogs by creating a bank of hypothetical excitatory frequency tuning curves for 172 auditory nerve fibers. The shape of each modeled tuning curve, plotted as a thin line in Fig. 5, was determined as a 4th order gammatone filter (*44-46*). The best sensitivity of each modeled tuning curve was adjusted to match the best excitatory frequency and threshold for a given nerve fiber based on previously published results reported in Fig. 1B of Ehret and Capranica (*31*). The bandwidth of each modeled tuning curve was estimated using the best-fit regression line for Q_10dB_ as a function of threshold [computed based on data from Fig. 2B in Ehret and Capranica (*31*)] to derive a predicted value of Q_10dB_ for each combination of best excitatory frequency and threshold. This estimate was used to compute the corresponding bandwidth 10 dB above threshold for each modeled tuning curve. (The joint relationship between best excitatory frequency, threshold, and bandwidth was not reported directly by Ehret and Capranica for (*31*) individual nerve fibers.)

### Social network analysis of calling survey data

To generate frog species co-calling networks, we utilized a publicly available dataset from the North American Amphibian Monitoring Program (NAAMP) (*38, 47*). Created as a citizen science collaboration between the United States Geological Survey (USGS) and a collection of state agencies, universities, and non-profit organizations, NAAMP was a long-term effort to monitor frog populations across 26 states in the eastern, central, and southern United States based on roadside calling surveys conducted by trained observers. Details about survey methods can be found elsewhere (*38, 47*). The NAAMP database consists of 319,765 observations of 57 identified species made during 21,934 roadside calling surveys conducted between 18 April 1994 and 9 August 2015. Fifteen of the 18 states encompassing the geographic range of green treefrogs, which are most abundant in the south-eastern United States, were included in the NAAMP dataset; therefore, coverage was high of geographic areas where green treefrogs were most likely to be heard calling. While the NAAMP dataset does not localize co-calling species precisely to the same body of water or close physical proximity, it represents the best (and only) measure of which other frog species might reasonably be expected to generate environmental noise for a green treefrog receiver in multi-species choruses across its geographic range on a continent-wide scale.

To create the co-calling network, we defined a node as a frog species and an edge as an event where both species were reported as calling during the same observation period (typically 3 min in duration) on the same date at the same survey stop at the same time. From networks containing only species co-occurring with green tree frogs (Figs. 6, S3), we identified the top ten co-callers by selecting those with the ten highest edge weights. We analyzed networks with and without the inclusion of any species complexes that were not resolved to individual species in the NAAMP dataset (e.g., *Pseudacris feriarum*/*fouquettei* complex). When using green treefrogs as the focal species, inclusion or exclusion of species complexes did not alter the top ten co-calling species identified. To aid in visualization of the network for Fig. 6, we took the square root of the raw edge weights and scaled this quantity by a factor of 15. All network analyses and images were generated in the *igraph* package in R (Version 3.3.2)(*48*).

### Statistical analyses

Unless indicated otherwise, a significance criterion of α = 0.05 was used. Two-tailed, one-sample t tests were used to test null hypotheses that there were no differences in tympanum vibration amplitude in response to free-field acoustic stimulation between the inflated and deflated states of lung inflation (*n* = 21). A two-tailed Pearson correlation was used to investigate the relationship between the peak frequency of the lungs in the inflated state and snout-to-vent length (*n* = 10). We used a two-tailed, paired-sample t test of the null hypothesis that deflating the lungs would not change the magnitude of the peak frequency of the lung’s resonance (*n* = 10). A two-tailed, paired-sample t test was used to directly compare the magnitudes of the resonance peak of the lungs in the inflated and reinflated states of inflation (*n* = 10). We also used two-tailed, one-sample t tests of the hypothesis that mean magnitudes of reduction in tympanum sensitivity were nonzero for three sound incidence angles (0°, -30°, and -60°) for each of five heterospecific species (15 comparisons total; *n* = 21 individuals for each comparison). All 15 comparisons remained significant after using the Holm-Šídák test to control for multiple comparisons; unadjusted P values are reported in Table S1.

## SUPPLEMENTARY MATERIALS

**Fig. S1.**
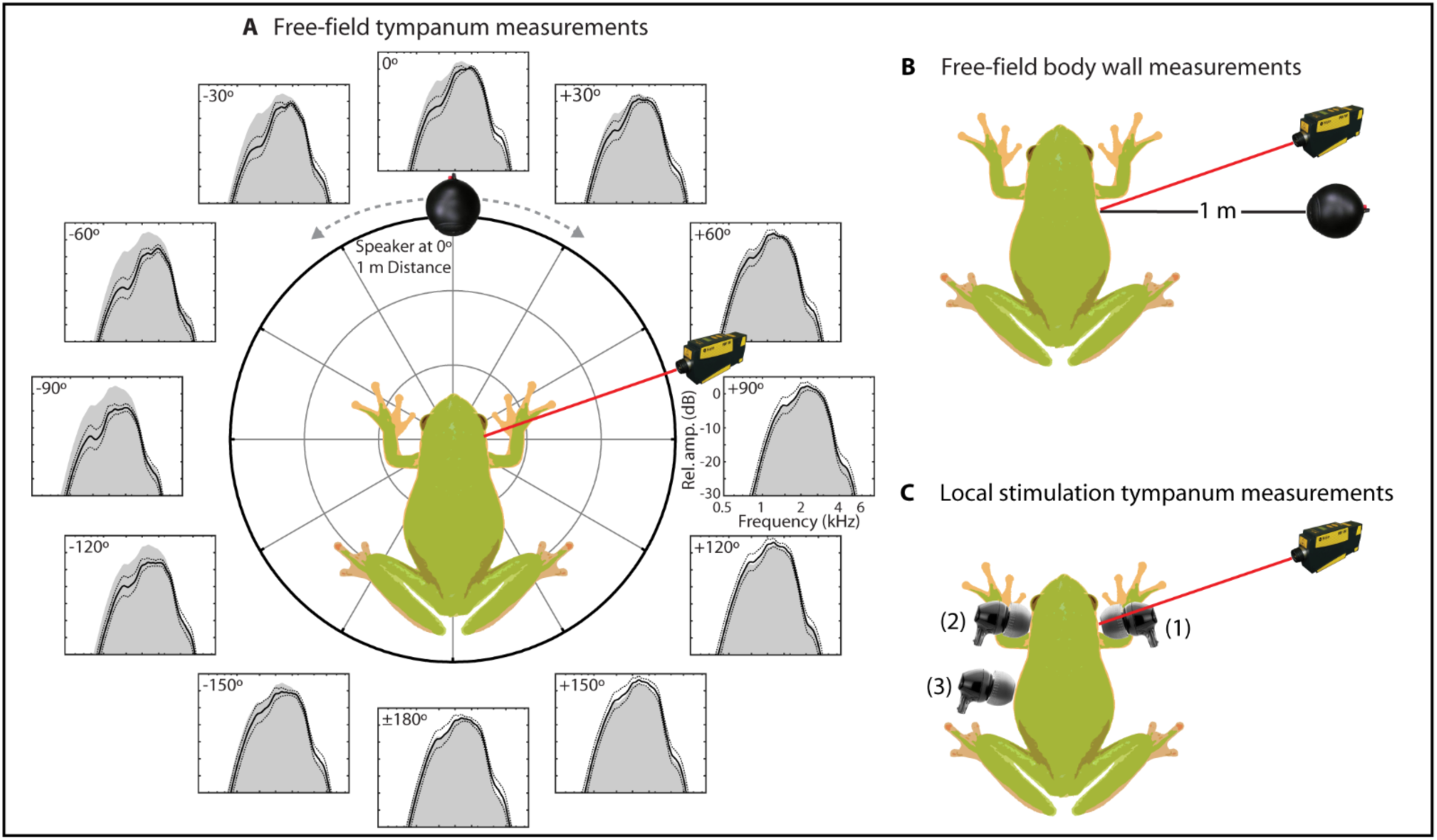
Laser vibrometry was used to measure the vibration amplitudes of the right tympanum and right body wall overlying the lung in response to acoustic stimulation. **(A)** The frequency spectrum of the vibration amplitude of the right tympanum was measured under free-field acoustic conditions in response to an FM sweep presented at each of 12 sound incidence angles (0° to ±180° in 30° steps) surrounding the animal from a speaker located 1 m away. These measurements were used to determine the impacts of the lungs on the tympanum’s response under conditions of free-field acoustic stimulation. *Insets*: Spectra show the tympanum’s mean (solid black line) ± 1 SD (dashed black lines) relative vibration amplitude as a function of frequency and sound incidence angle; the shaded gray area corresponds to the spectrum at +90° and is reproduced for comparison purposes at all other angles of sound incidence. **(B)** The frequency spectrum of the vibration amplitude of the right body wall overlying the lung was measured under free-field acoustic conditions in response to an FM sweep presented from a speaker located 1 m away at an ipsilateral (+90°) angle of sound incidence. These measurements were used to determine the resonance frequency of the lungs under conditions of free-field acoustic stimulation. (**C**) Local acoustic stimulation of the ipsilateral (to the laser) tympanum (1), the contralateral tympanum (2), and the contralateral body wall (3) was used to generate quantitative measures of the transmission gain of indirect sound input to the internal surface of the ipsilateral tympanum via the lungs and the contralateral tympanum. The laser used for measurements was always located at to the frog’s right side at an angle of +70° relative to the snout (at 0°).

**Fig. S2.**
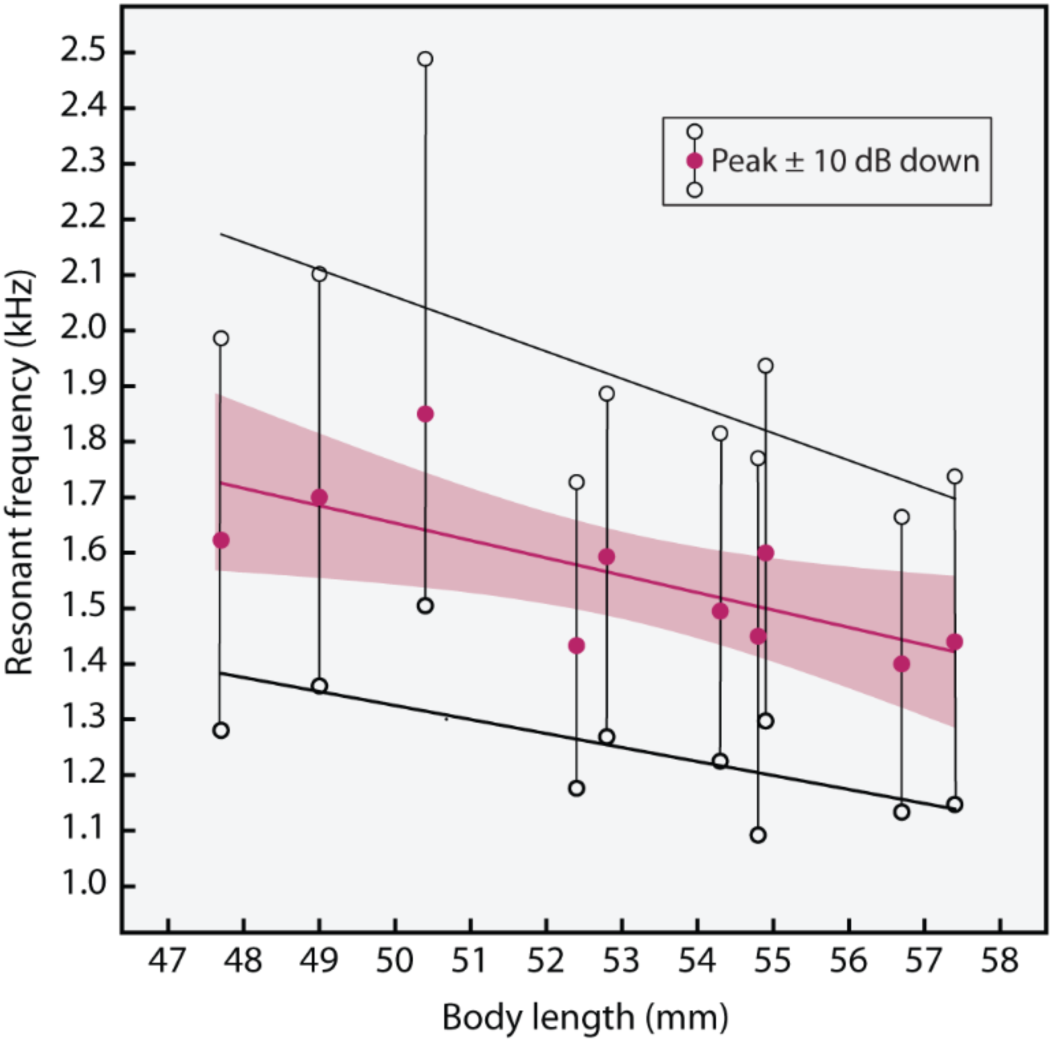
Frog lungs are size-dependent resonators. Under free-field acoustic stimulation by an FM sweep, the inflated lungs of green treefrogs exhibit a size-dependent resonance. Filled circles depict, as a function of body length, the mean peak resonance frequency of the body wall overlying the right lung in response to free-field acoustic stimulation by an FM sweep and measured using laser vibrometry. Open circles connected to each peak frequency represent the ± 10-dB down points on the resonance spectrum measured relative to the amplitude of the peak frequency. The colored line is the best-fit regression line (±95% CIs) for the peak frequency; black lines depict best-fit lines for the upper or lower 10-dB down points.

**Fig. S3.**
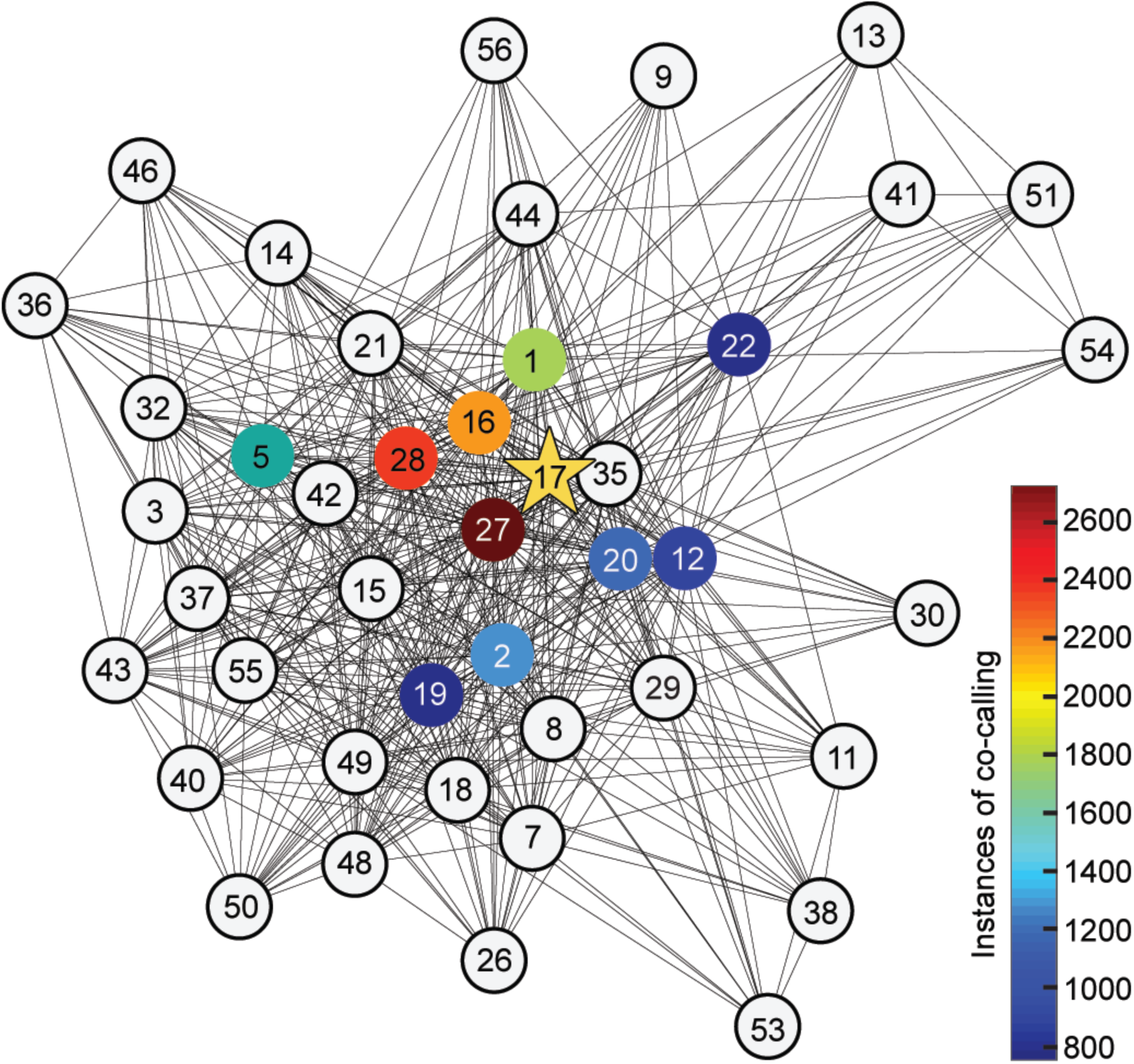
Social network analysis of data from the North American Amphibian Monitoring Program (NAAMP) identified 42 heterospecific, co-calling species. Using network analysis, we determined the incidence of “co-calling” between green treefrogs and other calling frog species in the NAAMP dataset. Green treefrogs were present on 9245 (2.9%) of the observations in the NAAMP dataset. Within these observations, a total of 42 heterospecific species co-called with green treefrogs. The top-ten co-callers are indicated by colored circles, with the number of observations of co-calling for each species (out of 19,809 total co-calling observations) depicted by the color map. The species identity for each numbered node is as follows: 1 *Acris crepitans*, 2 *Acris gryllus*, 3 *Anaxyrus americanus*, 5 *Anaxyrus fowleri*, 7 *Anaxyrus quercicus*, 8 *Anaxyrus terrestris*, 9 *Eleutherodactylus cystignathoides*, 11 *Eleutherodactylus planirostris*, 12 *Gastrophryne carolinensis*, 13 *Gastrophryne olivacea*, 14 *Hyla andersonii*, 15 *Hyla avivoca*, 16 *Hyla chrysoscelis*, 17 *Hyla cinerea*, 18 *Hyla femoralis*, 19 *Hyla gratiosa*, 20 *Hyla squirella*, 21 *Hyla versicolor*, 22 *Incilius nebulifer*, 26 *Lithobates capito*, 27 *Lithobates catesbeianus*, 28 *Lithobates clamitans*, 29 *Lithobates grylio*, 30 *Lithobates heckscheri*, 32 *Lithobates palustris*, 35 *Lithobates sphenocephalus*, 36 *Lithobates sylvaticus*, 37 *Lithobates virgatipes*, 38 *Osteopilus septentrionalis*, 40 *Pseudacris brimleyi*, 41 *Pseudacris clarkii*, 42 *Pseudacris crucifer*, 43 *Pseudacris feriarum*, 44 *Pseudacris fouquettei*, 46 *Pseudacris kalmi*, 48 *Pseudacris nigrita*, 49 *Pseudacris ocularis*, 50 *Pseudacris ornata*, 51 *Pseudacris streckeri*, 53 *Rhinella marina*, 54 *Scaphiopus couchii*, 55 *Scaphiopus holbrookii*, 56 *Scaphiopus hurterii*, 57 *Spea bombifrons*.

**Table S1.** Lung-mediated reductions in the tympanum’s response to frequencies in heterospecific calls.

| Species | Sound | Amplitude |  |  |  |
| --- | --- | --- | --- | --- | --- |
|  | incidence | reduction (dB) | *t* | *p* | df |
| *Lithobates clamitans* | 0° | 4.4 ± 2.0 | 4.38 | < 0.001 | 20 |
|  | -30° | 4.6 ± 2.2 | 3.98 | 0.001 | 20 |
|  | -60° | 5.4 ± 2.2 | 4.90 | < 0.001 | 20 |
| *Lithobates catesbeianus* | 0° | 4.0 ± 1.9 | 4.06 | 0.001 | 20 |
|  | -30° | 4.2 ± 2.2 | 3.70 | 0.001 | 20 |
|  | -60° | 5.0 ± 2.1 | 4.76 | < 0.001 | 20 |
| *Incilius nebulifer* | 0° | 4.0 ± 2.0 | 3.83 | 0.001 | 20 |
|  | -30° | 4.5 ± 2.6 | 3.43 | 0.003 | 20 |
|  | -60° | 3.8 ± 2.5 | 3.01 | 0.007 | 20 |
| *Anaxyrus fowleri* | 0° | 3.6 ± 1.9 | 3.63 | 0.002 | 20 |
|  | -30° | 3.8 ± 2.2 | 3.39 | 0.003 | 20 |
|  | -60° | 4.6 ± 2.1 | 4.43 | < 0.001 | 20 |
| *Hyla gratiosa* | 0° | 4.4 ± 2.0 | 4.38 | < 0.001 | 20 |
|  | -30° | 4.6 ± 2.2 | 3.98 | 0.001 | 20 |
|  | -60° | 5.4 ± 2.2 | 4.90 | < 0.001 | 20 |

## Acknowledgments

We thank C. Gerhardt and Marlene Zuk for helpful feedback on the manuscript; J. Tanner and M. Elson for help collecting frogs; and S. Gupta for assistance with animal husbandry; and Gary Calkins and the Texas Parks and Wildlife Division for permission to collect frogs under Scientific Permit Number SPR-0410-054

## Author contributions

N.L. collected and analyzed laser vibrometry data, wrote custom software for acquiring laser vibrometry data, conducted the bioacoustics analyses, and co-wrote the manuscript; J.C-D. collected and analyzed data and edited the manuscript, L.A.W. conducted the social network analysis, provided relevant text, and edited the manuscript; K.M.S. made recordings of frogs in the field and edited the manuscript; and M.A.B. conceived and led the project, secured funding, collected and analyzed data, and co-wrote the manuscript

## Data and materials availability

The datasets generated and analyzed during the current study will be made available before publication in the Data Repository for the University of Minnesota (https://conservancy.umn.edu/handle/11299/166578) and are available to reviewers and editors upon request from the corresponding author. All code and data used to generate the social network figures are available at: https://github.com/whit1951/FrogNetworks.

